# Self-Attention based model for *de-novo* antibiotic resistant gene classification with enhanced reliability for Out of Distribution data detection

**DOI:** 10.1101/543272

**Authors:** Md-Nafiz Hamid, Iddo Friedberg

## Abstract

Antibiotic resistance monitoring is of paramount importance in the face of this ongoing global epidemic. Using traditional alignment based methods to detect antibiotic resistant genes results in huge number of false negatives. In this paper, we introduce a deep learning model based on a self-attention architecture that can classify antibiotic resistant genes into correct classes with high precision and recall by just using protein sequences as input. Additionally, deep learning models trained with traditional optimization algorithms (e.g. Adam, SGD) provide poor posterior estimates when tested against Out-of-Distribution (OoD) antibiotic resistant/non-resistant genes. We train our model with an optimization method called Preconditioned Stochastic Gradient Langevin Dynamics (pSGLD) which provides reliable uncertainty estimates when tested against OoD data.

## 1 Introduction

Antibiotic resistance is a global scourge that is taking an increasing toll in mortality and morbidity, in both nosocomial and community acquired infections[1,2]. A growing number of once easily treatable infectious diseases such as tuberculosis, gonorrhea, and pneumonia are becoming harder to treat as the scope of effective drugs is shrinking. The CDC estimates that 2,000,000 illnesses and 23,000 people die annually from antibiotic resistance in the US alone[3]. The overuse of antibiotics in health care and agriculture is exacerbating the problem to the point that the World Health Organization is considering antibiotic resistance “one of the biggest threats to global health, food security and human development today”. Identifying genes associated with antibiotic resistance is an important first step towards dealing with the problem[4], and providing a narrow-spectrum treatment, targeted solely against the types of resistance displayed. This statement is especially true when dealing with genes acquired from human or environmental metagenomic samples[5]. A rapid identification of the class of antibiotic resistance that may exist in a given environmental or clinical microbiome sample can provide immediate guidance to treatment and prevention.

In this study, we developed a deep neural network that can predict antibiotic resistant genes into 15 classes from protein sequences as well as detect OoD data - genes resistant against antibiotic classes that were not included during training, or non-antibiotic resistant genes. This property can be useful in identifying metagenomic sample resistance for the purpose of providing a focused drug treatment. Traditional methods [6, 7, 8] to identify antibiotic-resistant genes usually take an alignment based best-hit approach which causes the methods to produce many false negatives [9]. Recently, a deep learning based approach was developed that used normalized bit scores as features that were acquired after aligning against known antibiotic resistant genes [9]. In contrast, our model only uses the raw protein sequence as its input, and as such doesn’t need do any kind of alignment against a set database.

At the same time, neural networks are known for providing high confidence scores on inputs that are from a different probability distribution than the model was trained on [10, 11]. This can result in disastrous consequences in sensitive applications such as health care or self-driving systems. Here, we develop deep learning models trained with Preconditioned Stochastic Gradient Langevin Dynamics (pSGLD) [12] as well as a traditional optimization method known as ADAM [13]. Both models give significant accuracy on the test set in terms of predicting antibiotic resistance solely from the protein sequence. But we show that the model trained with pSGLD is better equipped to predict OoD data i.e., it assigns a low probability to sequences from proteins that are not related to antibiotic-resistance or are from classes that were not included in training.

## 2 Detection of Out-of-distribution data with Preconditioned Stochastic Gradient Langevin Dynamics (pSGLD)

Typically, neural networks are trained with optimization methods such as Stochastic gradient descent (SGD) [14] or its variants such as Adam [13], Adagrad [15], RMSprop [16] etc. In SGD, for each iteration a mini-batch from the dataset is used to update the parameters of the neural network. For each iteration *t*, training data *X_t_* = {*x*_*t*1_,…,*x_tn_*} is provided, and for parameters *θ*, the update Δ*θ_t_* is:

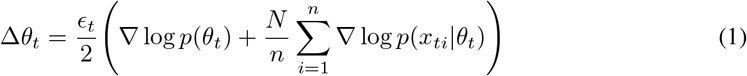

At the same time, SGD or its variants do not capture parameter uncertainty. In contrast, Bayesian approaches such as Markov Chain Monte Carlo (MCMC) [17] techniques do capture uncertainty estimates. One such class of techniques are Langevin dynamics [18] which inject Gaussian noise into Equation 1 so that the parameters do not collapse into the Maximum a posteriori (MAP) solution:

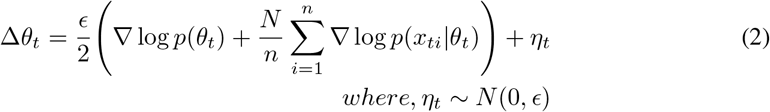

However, MCMC techniques require that the algorithm go over the entire dataset per iteration before making a parameter update. This slows down the model training process, and also requires huge computational costs. To remove this problem, Stochastic Gradient Langevin Dynamics (SGLD) was introduced [19], which combined the best of both worlds i.e. inserting Gaussian noise into each mini-batch of training data. In SGLD, during each iteration for SGD, Gaussian noise is injected which has a variance of the step-size *ϵ_t_*:

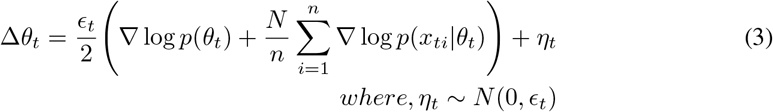

This injection of Gaussian noise has an advantageous side-effect, as it also provides a better calibration of confidence scores of predictions on OoD data. For example, [10] showed that an SGLD trained neural network provides low confidence scores when trained on the MNIST [20] dataset but tested on the NotMNIST dataset [21]; whereas an SGD trained neural network still naively provides high confidence scores. We used preconditioned SGLD (pSGLD) [12] - a variation of SGLD where we introduce noise to RMSprop, a modificaiton of the SGD algorithm - to train a neural network to classify protein sequences into their antibiotic resistance classes. In the experiment section, we show that a pSGLD trained network provides really high accuracy when classifying antibiotic resistant genes for the classes it was trained on while providing low confidence scores when doing prediction on OoD protein sequences. In contrast, an ADAM trained model provides high accuracy when classifying antibiotic resistant genes for the classes it was trained on as well as when doing prediction on OoD protein sequences.

## 3 Experimental setup

### Dataset

We used a modified version of the dataset curated in the DeepARG study [9]. Briefly, The dataset was created from the CARD [22], ARDB [23] and UNIPROT [24] databases with a combination of computational and manual curation. The original dataset has 14974 protein sequences that are resistant to 34 different antibiotics (our classes in the multi-class classification task). There were 19 classes that had training samples of 11 sequences or less. We discarded these classes and were left with 15 classes with a total of 14907 protein sequences.

### Model

We used a self-attention based sentence embedding model introduced in [26]. Figure 1 shows a schematic of the architecture of the model we used for antibiotic resistant gene classification. For input, we represented each amino acid in a protein sequence as an embedding vector which was randomly initialized, and then trained end-to-end. We used Long short term memory (LSTM) [25] recurrent neural network which takes these embedding vectors as input. Following that is the self-attention part which we can think of as a feed-forward neural network with one hidden layer. This network takes the output from the LSTM layer as input, and outputs weights which are also called attentions. We multiplied the outputs of the LSTM layer with these attentions to get a weighted view of the LSTM hidden states. This multiplication result gets passed onto the next layer of the network, and the final layer is a softmax layer which assigns a probability to each of the 15 classes of antibiotics that add upto 1.

**Figure 1:**
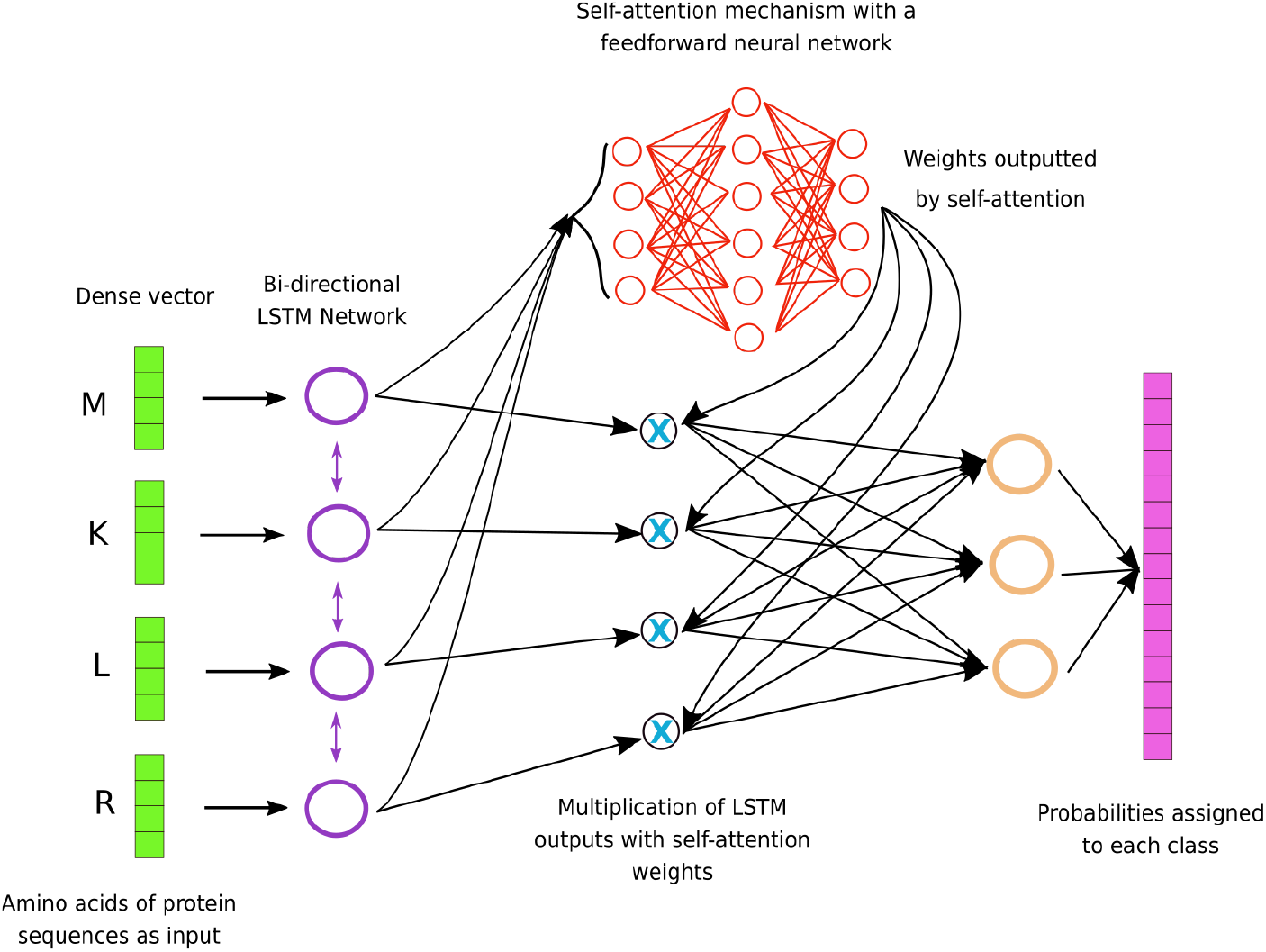
Our self-attention model used for antibiotic resistant gene classification. Each amino acid in a protein sequence is represented by a vector. These vectors are used as input for LSTM neural networks [25]. Outputs of the LSTM network goes into a simple feed-forward network, and the output of this feed-forward network are weights which sum upto 1. These weights are multiplied with the outputs of the LSTM network to give them weights. This mechanism lets the model learn which input from the protein sequence is crucial in classifying a gene into its correct antibiotic class. Finally, there is a softmax layer, the size of which is 15 corresponding to the 15 antibiotic classes we have. This layer assigns a probability to each class, the class that is assigned the highest probability becomes the prediction for a certain protein sequence.

The model that is trained with Adam has one bi-directional LSTM layer of size 64. The embeddings for each amino acid are of size 10. Dropout of 0.4 was used for regularization. We used a learning rate of 0.001 with a weight decay value of 0.0001 for ADAM. For pSGLD, the model has one bi-directional LSTM layer of size 128. Embedding vectors for amino acids are of size 400. A learning rate of 0.001, and dropout of 0.4 was also used for pSGLD.

We divided our dataset into a 70/20/10% training, validation, and test set split. We trained our model with Adam and pSGLD on the training dataset, and tuned the hyper-parameters by checking the performance on the validation dataset. We report both models’ performances on the test set.

## 4 Results

Figure 2 shows the performance of Adam and pSGLD trained models on the test set in terms of Precision, Recall and *F*_1_ for each class and overall.

**Figure 2:**
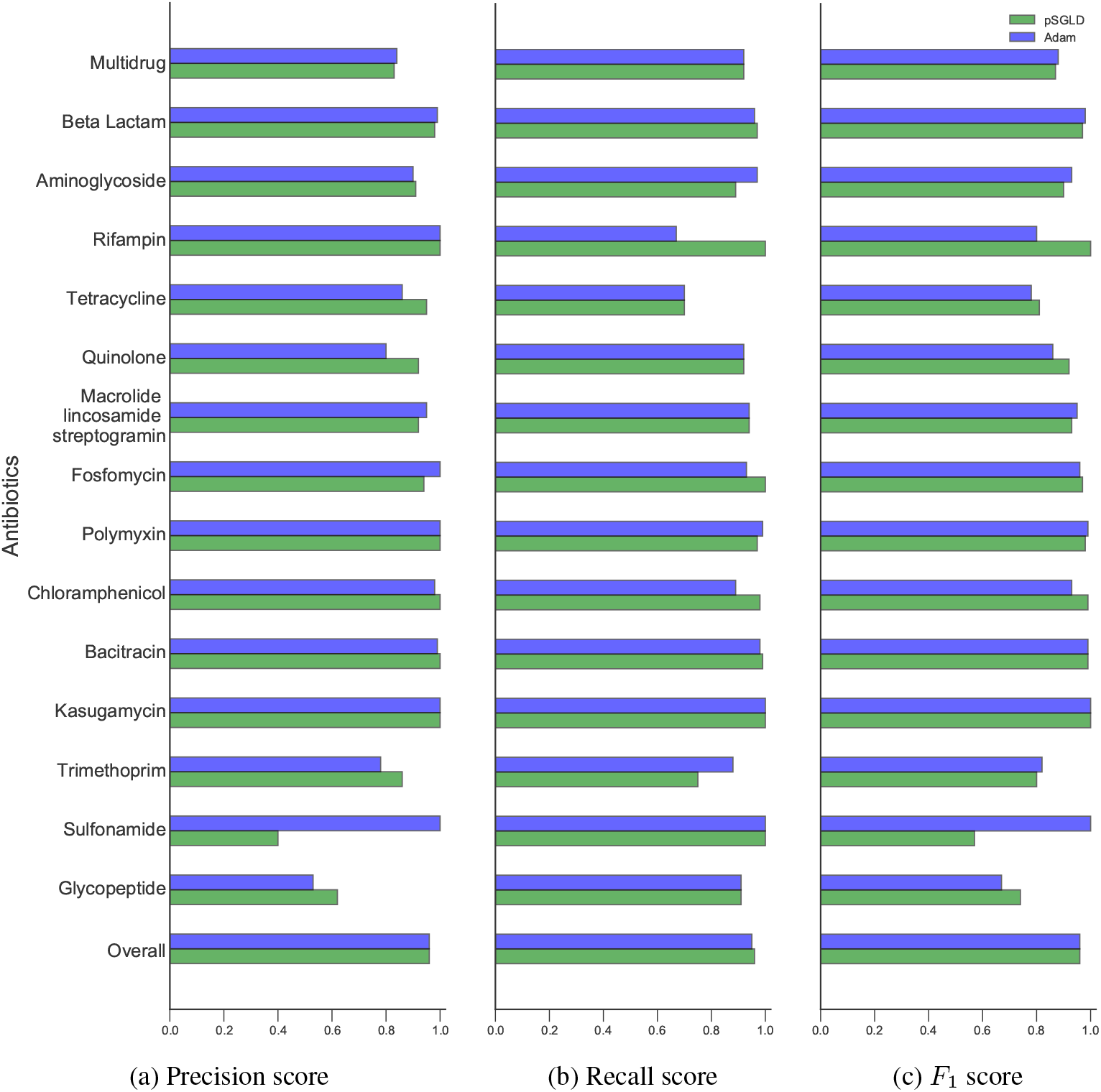
Precision, Recall, and *F*_1_ score for Adam and pSGLD. Our deep learning models have really high accuracy for antibiotic resistant gene classification without using any kind of alignment, and just using the amino acids in protein sequences as features.

Precision (*Pr*), Recall (*Rc*), and *F*_1_ are defined as:

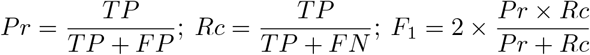

Where *TP*: True Positives, *FP*: False Positives, *FN*: False Negatives.

We can see that overall the performance of both models are comparable. Adam has a 96% precision, 95% recall, and 96% *F*_1_ score whereas pSGLD has a 96% precision, 96% recall, and 96% *F*_1_ score.

Figure 3 shows a histogram distribution of probability assigned to a class predicted for a protein sequence by both models. Figure 3a shows the distribution of probability of a predicted class when both models are correct. Figure 3b shows the distribution of probability of a predicted class when both models are incorrect. We can see that when the models are predicting correctly, the probability assigned to the correct class by both models are high. In contrast, when the models are incorrect, pSGLD tends to assign somewhat lower probability to the predicted class signalling its uncertainty.

**Figure 3:**
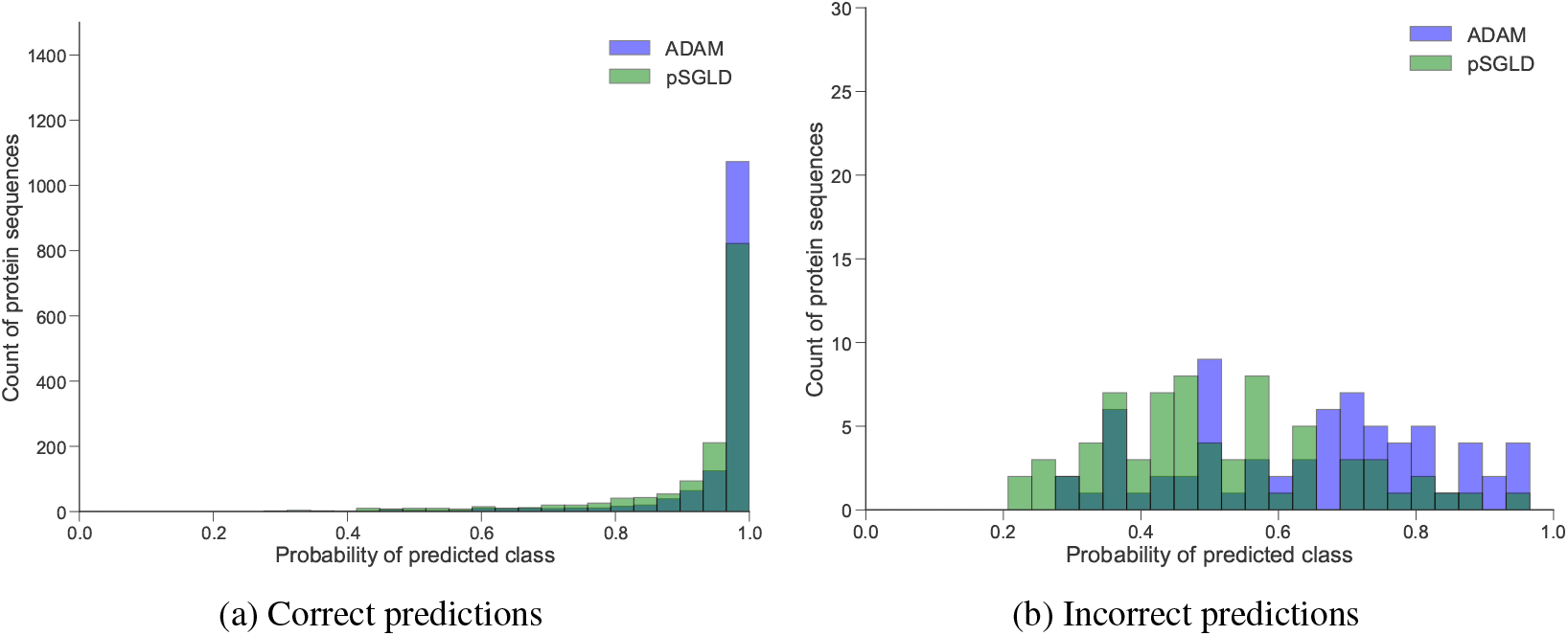
Probability assigned to the class predicted by the pSGLD, and ADAM models. (**a**) Probability assigned to a class when the models are correctly predicting. (**b**) Probability assigned to a class when the models are incorrectly predicting. Scale of the y-axis in (a) and (b) are different for ease in visibility.

### Result on OoD data

In the next step, we tested both models on OoD samples. For this we used three datasets - **(i)** The 19 classes out of the 34 antibiotic resistant classes from the DeepARG dataset that we discarded. We did not include protein sequences from these classes in training and testing our models. These 19 classes have a total of 67 protein sequences. **(ii)** 19,576 human genes collected from UNIPROT [27] that we can confidently assume are not antibiotic resistant. **(iii)** 480 bacteria genes that are involved in the metabolism pathway were curated from the KEGG [28] database. These have very low chances of involvement in antibiotic resistance.

Before testing on these sequences, our expectation is that an ideal model trained on a different set of classes (the 15 antibiotic classes used in training and testing in our case) should provide low probabilities for its prediction on out-of-class sequences. The model should also provide low probabilities for Human genes and Bacteria metabolism genes that are not antibiotic resistant.

Figure 4 shows a histogram distribution of probability assigned to the predicted class for both models on these sequences. We can see from the figure that in all three cases the probability distribution for pSGLD is centered around 0.5 or left whereas for ADAM the distribution is heavily right skewed. The ADAM trained model is still predicting these OoD sequences to be one of the 15 classes it was trained on with high confidence. In contrast, the pSGLD model is conveying its uncertainty over its predictions.

**Figure 4:**
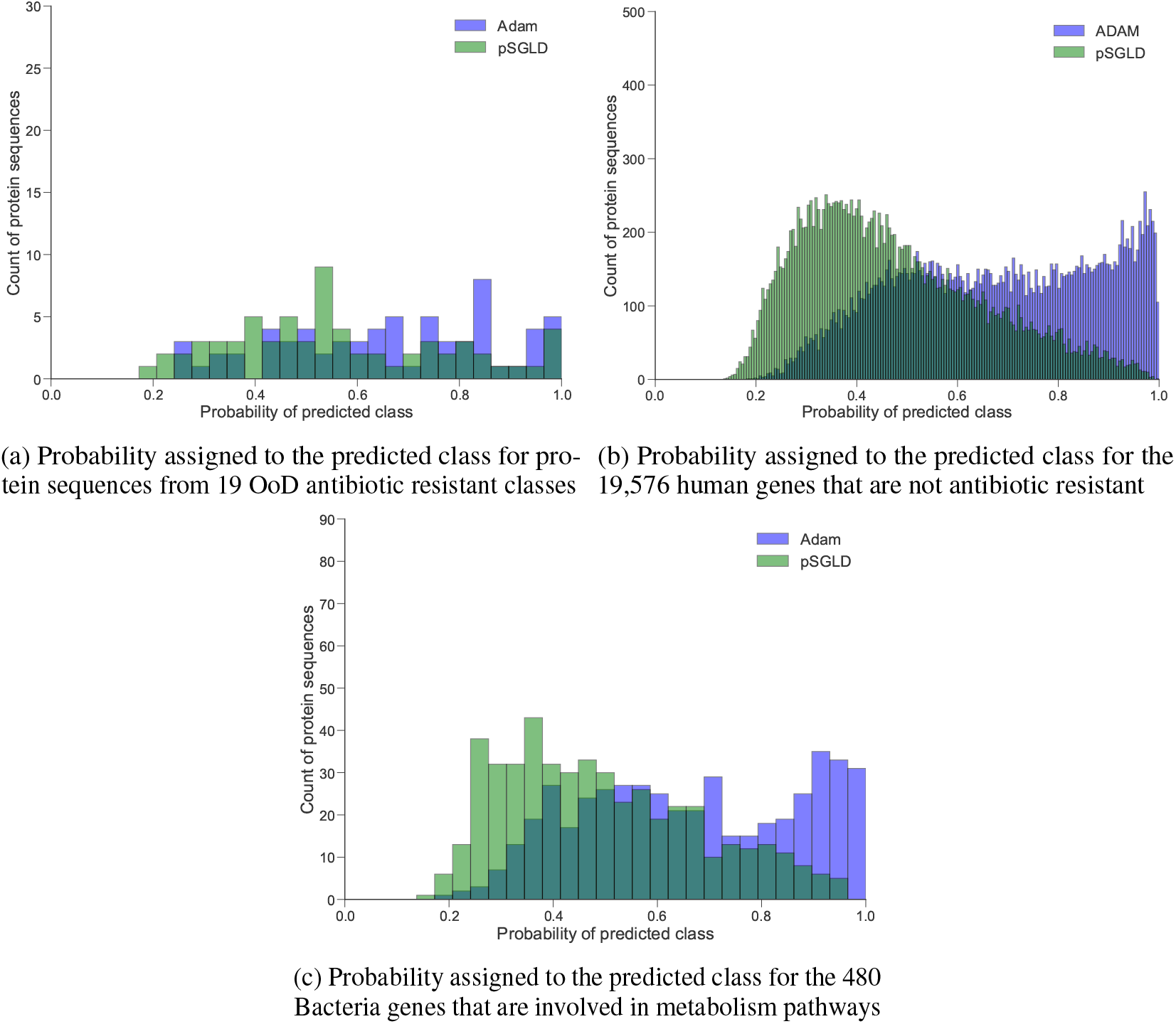
Probabilities assigned to the predicted class by both pSGLD and ADAM trained models. The pSGLD trained model has left-skewed probability distribution on the OoD protein sequences while the ADAM trained model has right-skewed probability distribution implying high confidence in its predictions. Scale of the y-axis in (a), (b), and (c) are different for ease in visibility.

Next, we tested both models in a binary classification setting where our positive class includes the test set of the 15-class dataset from DeepARG. This test set has 1491 genes. For our negative dataset, we used the three OoD datasets mentioned previously. We tested both models, and measured their performance in terms of classifying the negative class from the positive class. The models were not trained to do this binary classification. They were still only trained to predict an antibiotic class from the 15 classes. But we are testing them to see how good they are when facing negative data just like in a real world scenario. This type of training to do a different task but then testing to detect OoD data is also used in computer vision to measure the performance of a system on identifying OoD data [29, 30].

Figure 5 shows the Precision-Recall (PR) curves and Receiver Operating Characteristic (ROC) curves for the binary classification task. We can see that pSGLD model has PR and ROC curves that cover a greater area than the ADAM model when negative data are the Humand and Bacteria genes. Specially, for Human genes, performance of pSGLD is significantly better implying pSGLD can better detect data that it has not seen in training. pSGLD also performs better when negative data is Bacteria genes that are not involved in antibiotic resistance. Performance of pSGLD and ADAM is almost equal when tested on the 67 genes from antibiotic resistant classes that were not included in the training dataset.

**Figure 5:**
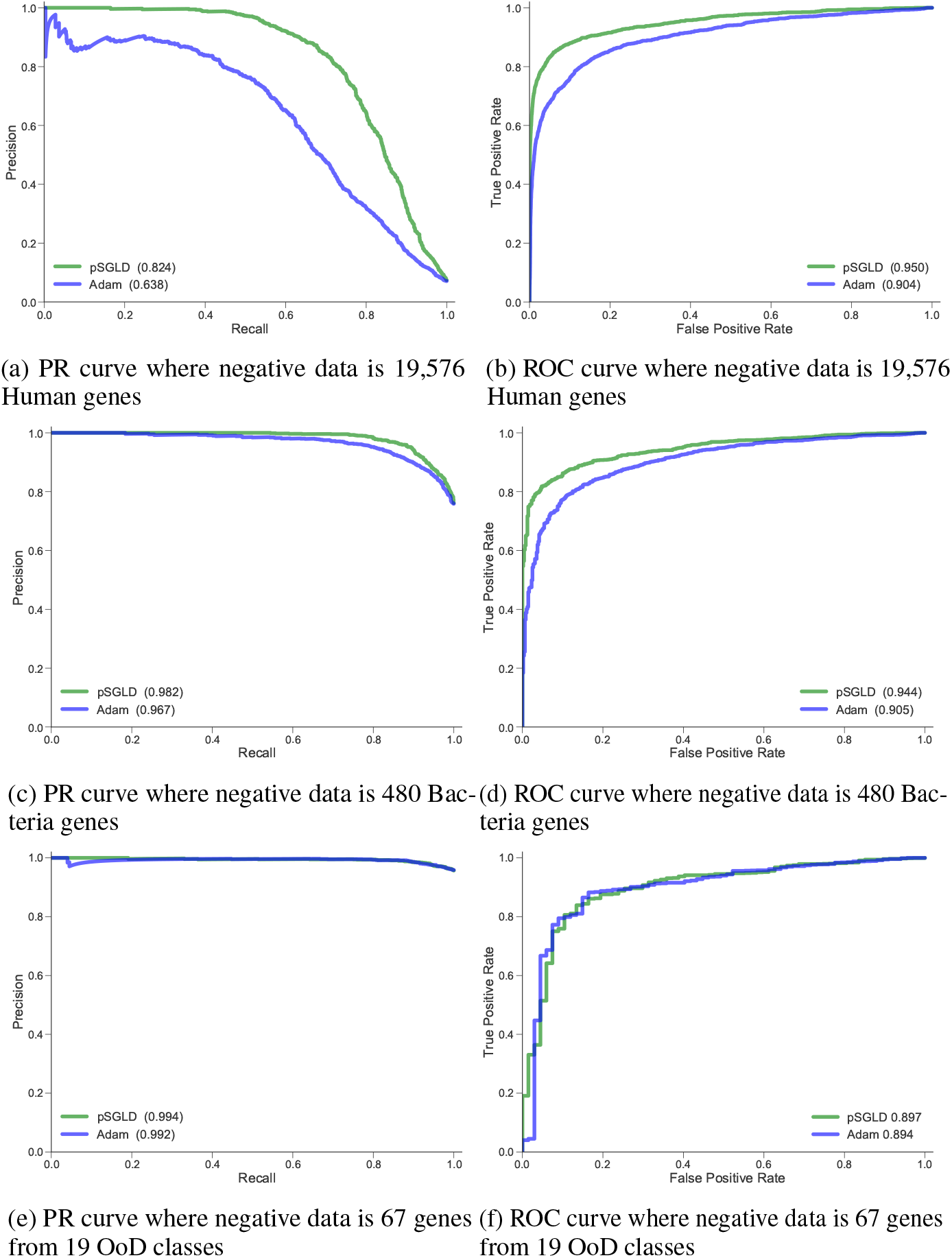
Precision-Recall (PR) curves and Receiver Operating Characteristic (ROC) curves for our binary classification task where for all cases positive data consists of the test set from our modified DeepARG dataset. These are 1491 genes that belong to one of the 15 antibiotic classes from the modified DeepARG dataset. For **a** and **b**, negative data is 19,576 Human genes that do not possess antibiotic resistance. For **c** and **d**, negative data is 480 Bacteria genes involved in the metabolism pathway, and do not possess antibiotic resistance. For **e** and **f**, negative data is 67 genes that are antibiotic resistant to 19 classes that were not included in the training and testing of our models.

## 5 Discussion

In summary, the contributions of this paper are as follows:

1. The self-attention based deep learning model provides extremely high accuracy in terms of precision, recall, and *F*_1_ for antibiotic resistant gene classification. This is of crucial importance in many clinical settings. Additionally, the model only uses raw protein sequences as its input removing any necessity for additional feature processing. In contrast, the DeepARG [9] method first creates a similarity distance matrix by aligning sequences against CARD and ARDB datasets. This similarity matrix is used as features for the deep neural network.
2. The self-attention based deep learning model trained with pSGLD provides reliable confidence scores for OoD gene detection. In the real world, we cannot expect that a model will see only antibiotic resistant genes that belong to one of the antibiotic classes that were included in its training. The model may encounter antibiotic resistant genes that are resistant to antibiotics not included in its training set, or genes that are not antibiotic resistant at all. We need the model to provide low confidence scores when it encounters one of these two scenarios.
3. *De-novo* antibiotic resistant gene classification overcomes the weakness of lots of false negatives resulting from alignment based methods such as BLAST [31]. This can be seen from the high recall score of our models. We refer to the DeepARG [9] paper for a detailed expound on the weakness of predicting a high number of false negatives by BLAST.
4. With the self-attention based model, we present a viable use case of using entity embeddings [32] as inputs for categorical variables. Amino acids are categorical variables, and usually represented in machine learning by one-hot vectors - vectors of size 20 where all indexes are zeros except the index of the amino acid we are representing which is 1. We show that, instead of using a one-hot vector, if we use a dense vector - a vector which has all non-zero real values - that is randomly initialized at the beginning, and then trained end-to-end inside the neural network, we can gain significant stride in terms of accuracy. These are known as entity embeddings in the literature. Also, there has been a trend on using word2vec type unsupervised learning to learn dense vectors for protein *k*-mers in biology [33]. Our results show that there is a need to consider end-to-end learning of dense vectors like in our model which can save precious time and computing that is needed for unsupervised learning.
5. DeepARG is based upon the Theano [34] deep learning framework which has been deprecated recently, and no longer being developed. This makes it difficult for new users to use the software. In contrast, our model is developed with the Pytorch [35] deep learning framework which is being actively developed, and is used by a huge number of deep learning researchers. This will make it easier for new users to adopt the software which will be released soon.

In this study, we applied a self-attention based deep learning model to a multi-class classification task of antibiotic resistance type classification from protein sequences. We trained our model with two different optimization algorithms, ADAM and pSGLD. The overall *F*_1_ score for both the ADAM and pSGLD trained models are 96%. Both models predict the antibiotic class of resistant genes with high accuracy. Comparison with the DeepARG model was not feasible as they use the CARD and ARDB protein sequences for generating features by aligning them against the UNIPROT protein sequences they curated. We could not find out, specifically, which protein sequences they used as their test data. Our model used all of the protein sequences from CARD, ARDB, and UNIPROT, and divided them into training, validation, and test set after shuffling them. Moreover, the test set performance reported in the DeepARG model suffers from data leakage. They selected protein sequences from UNIPROT by using an alignment threshold against the CARD and ARDB dataset when creating the complete dataset. Then they aligned the same selected UNIPROT sequences against CARD and ARDB, and used the bit scores to make a similarity distance matrix. This matrix was used as features for their neural network to predict on test data that were previously selected from UNIPROT by alignment threshold against CARD and ARDB.

We also tested both of our models on three datasets of protein sequences that we know either belong to classes of antibiotic resistance that were not part of our training and testing, or are not antibiotic resistance associated. The ADAM trained model predicted them to be of the 15 classes with high confidence scores. In contrast, the pSGLD trained model provided predictions with a lower probability distribution for the proteins not associated with antibiotic resistance. We hypothesize that the Gaussian noise introduced in the pSGLD training scheme impedes the neural networks to completely collapse on the Maximum Likelihood solution. Therefore, a pSGLD trained model lets a discriminative model detect OoD data points, and provide lower probabilities in its predictions for them. This is an important property, especially when we consider the open world problem in biology where for any classification task it is hard to collect negative training samples for training the machine learning algorithm [36]. Future research might involve improving our neural network training methods so that the probability distribution of prediction for in-distribution and out-of-distribution data has much more stark difference than what we could manage with pSGLD. For that, we can take inspiration from the research on detection of adversarial instances [37] done by the computer vision community.

